# Distinct spread of DNA and RNA viruses among mammals amid prominent role of domestic species

**DOI:** 10.1101/421511

**Authors:** Konstans Wells, Serge Morand, Maya Wardeh, Matthew Baylis

**Affiliations:** Department of Biosciences, Swansea University, Swansea SA2 8PP, Wales, UK; CIRAD ASTRE, CNRS ISEM, Faculty of Veterinary Technology, Kasetsart University, Bangkok, Thailand; Department of Epidemiology and Population Health, Institute of Infection and Global Health, University of Liverpool, Leahurst Campus, Chester High Road, Neston, Cheshire CH64 7TE, UK; Health Protection Research Unit in Emerging and Zoonotic Infections, University of Liverpool, UK

**Keywords:** cross-species transmission, disease emergence, disease risk assessment, host-parasite interaction, pathogen spillover, zoonotic disease risk, network analysis, virus spread

## Abstract

Emerging infectious diseases arising from pathogen spillover from mammals to humans comprise a substantial health threat. Tracing virus origin and predicting the most likely host species for future spillover events are major objectives in One Health disciplines. However, the species that share pathogens most widely with other mammals, and the role of different wildlife groups in sharing viruses with humans remain poorly identified. To address this challenge, we applied network analysis and Bayesian hierarchical models to a global database of mammal-virus associations. We show that domesticated mammals and some primates hold the most central positions in networks of known mammal-virus associations. We revealed strong evidence that DNA viruses were phylogenetically more host specific than RNA viruses, while the frequencies of sharing viruses among hosts and the proportion of zoonotic viruses in hosts were larger for RNA than DNA viruses. Among entire host-virus networks, Carnivora and Chiroptera hold central positions for mainly sharing RNA viruses with other host species, while network centrality of Primates scored relatively high for sharing DNA viruses. Ungulates hold central positions for sharing both RNA and DNA viruses. Acknowledging the role of domestic species in addition to host and virus traits in patterns of virus sharing is necessary to improve our understanding of virus spread and spillover in times of global change.

## 1. Introduction

Pathogen spillover and cross-species transmission between animals and humans is a major source of infectious diseases and a considerable global public health burden [1, 2]. Understanding the factors that enable or facilitate these processes is a crucial step for such events to be predicted. Host shifting, that is the colonization of a new host species by a pathogen, requires a certain level of overlap in species traits (‘ecological fitting’) in order to overcome barriers of cross-species transmission and for survival and reproduction within novel host species [3-5]. In the search for mechanisms and enabling conditions that may help to predict the future emergence of infectious diseases from animal populations, the necessity of considering entire host species communities amongst underpinning biogeographic structure and connectivity have been recently emphasized [6-9].

Network analyses that describe the connections of different host species in terms of parasite sharing have proven useful in analysing host specificity and parasite spread [10, 11], particularly since they offer the opportunity to explore community-wide pathogen spread (the distribution of a pathogen among host species, a pattern emerging from past and contemporary host shifting events that connect host species as nodes in a network). Such approaches may lead to increased predictability of future pandemics; for example, some recent “big data” studies of mammal-virus associations have explored whether host traits and geographic distribution can predict those species to most likely harbour undiscovered viruses that may cause future pandemics [11-13].

Among different host groups, bats have come into particular focus for harbouring zoonotic viruses (viruses detected in humans and at least one other mammal species) as pandemics of Nipah, SARS and Ebola were found to be linked to bats as major reservoirs between 2001 and 2005 [14]. Yet despite these important advances in virus discovery and analytical approaches, our understanding of virus sharing and their spread through entire networks of mammalian host species remains limited. The challenge of assessing different animal species in their role for virus spread is understandable, as detailed information about virus sharing across entire communities became only recently available [12, 15] amid the challenge that many virus species remain unknown [16].

We address this knowledge gap by exploring the role of different mammalian species in the spread of viruses through entire host communities. In particular, we tested whether domestic species (livestock and companion animals) play a major role in virus spread and spillover among humans and wildlife. To this end, there are strong reasons why domesticated animals should cover central positions in networks of host-virus associations. Domesticated animals share large numbers of viruses and other parasites with humans [17] and were recently reported to play crucial roles in the sharing of helminth parasites between humans and wildlife [9]. Moreover, the large numbers of domestic animals compared to those of wildlife [18], and close contact between them and people, creates ground for frequent and multilateral exposure. For entire networks of viruses and mammalian host associations, we also expect that virus sharing differ between the two different genome types of DNA and RNA viruses. Faster replication and higher genetic diversity in RNA viruses have been proposed to likely increases their host range through more frequent host shifting and adaptation to distantly related host species, whereas DNA viruses and retroviruses are assumed to be more host-specific due to codivergence with their hosts over much longer evolutionary timescales [19-22]. Yet, to date little comprehensive work has been conducted of whether host sharing and spread at the network level differ among these viruses and whether they interact with the various groups of mammals in different ways. We used network centrality analysis and Bayesian hierarchical models to quantify the extent of virus sharing among different mammalian host species and the proportion of zoonotic viruses carried in different hosts. Within the hierarchical model of virus sharing, probabilistic estimates of the variation in the pairwise phylogenetic and functional similarities of infected and potential host species provides a signal of host specificity.

## 2. Material and methods

### (a) Virus-host data

We extracted mammal-virus species-level interactions from the Enhanced Infectious Diseases Database (EID2) [15] in the version from May 2018. In brief, EID2 utilises automated procedure to extract information on pathogens, their hosts and locations from two sources: 1) the meta-data accompanying nucleotide sequences published in the National Center for Biotechnology Information (NCBI) Nucleotide database (www.ncbi.nlm.nih.gov/nuccore); and 2) titles and abstracts of publications indexed in the PubMed database (www.ncbi.nlm.nih.gov/pubmed). To date, EID2 has extracted information from 7,1076,379 sequences (and processed 100M+ sequences), and 8,643,203 titles and abstracts. EID2 imports names of organisms (1,407,577 of which 157,894 are viruses and 10,071 are mammals) and their taxonomic hierarchy from the NCBI Taxonomy database (http://www.ncbi.nlm.nih.gov/Taxonomy/), and supplements it with an exhaustive collection of alternative names. EID2 also contains information on 2,171 organisms of interest that were not found in the NCBI Taxonomy database (none of which were mammals or viruses). In general, EID2 follows the NCBI definitions of ‘species’, ‘subspecies’ and ‘no rank’, with most unclassified and uncultured species are denoted ‘no rank’. After applying the classification rules listed in [15], EID2 contains 15,959 species of viruses and 6.735 species of mammals.

The data of interest for this study were associations of mammalian (including human) species with different virus species, independent of location records. We considered a mammalian species to be host to a virus if at least four independent publications or one sequence meta-data reported an association between the host (or any of its subspecies) and the virus (or any of its subspecies or strains). Virus species were assigned to genome type (DNA, RNA or other/unspecified) following NCBI taxonomy as utilised by EID2.

Mammal species synonyms and taxonomic orders were standardized using the taxonomy of [23], the online version of IUCN Red List and Integrated Taxonomic Information System, ITIS (accessed May 2018). This revision enabled us to match the most recent host names to trait data.

Of the 816 non-human mammalian host species in our data set, we considered 20 species as ‘domestic’ (including the major commensal rodent species) and all other as ‘wildlife’. Domestic species were banteng (*Bos javanicus*), yak (*B. mutus*), cow (*B. taurus*), bactrian camel (*Camelus bactrianus* and *C. ferus*), dromedary (*C. dromedarius*), dog (*Canis familiaris* and *Canis lupus*), goat (*Capra aegagrus*), guinea pig (*Cavia porcellus*), wild ass (*Equus africanus*), donkey (*E. asinus*), horse (*E. caballus*), cat (*Felis catus*), guanaco (*Lama guanicoe*), house mouse (*Mus musculus*), rabbit (*Oryctolagus cuniculus*), sheep (*Ovis aries*), brown rat (*Rattus norvegicus*), black rat (*R. rattus*), pig (*Sus scrofa*) and vicugna (*Vicugna vicugna*). We constrained our domestic species selection to these major domestic species only to showcase possible differences in pathogen sharing, while we are aware that there are some additional species that may be considered to be domestic animals.

We generated four different measures of sampling effort for each mammalian host species, namely 1) number of publications (summed over all associated virus species), 2) number of virus sequences recorded (summed over all associated virus species), 3) Shannon diversity of publication records, accounting for the proportional number of publications for each associated virus species and 4) Shannon diversity of sequence records, accounting for the proportional numbers of sequence records for each associated virus species. For Shannon indices larger values are linked to overall larger number of records and a more even distribution of records among different virus species, i.e. higher overall sampling coverage [24]. We generated these multiple indices as proxies of sampling intensity, as the true sampling effort is not known. This is because records of species interactions in the literature are arguable ‘presence-only’ records and rarely report the lack of interactions that would reduce the number of pseudo-absences in biotic interaction data [25, 26].

### (b) Mammalian host phylogeny and ecological trait data

A goal of this study was to assess whether variation in the phylogenetic and ecological similarities of mammalian species predict patterns of virus sharing (i.e. pairs of mammalian specie infected by the same virus species; the entire interaction matrix of all mammal-virus interactions constitutes the backbone of the ecological network of how mammalian species are connected in terms of virus sharing) and the proportion of zoonotic viruses associated with different host species. We gathered ecological trait data from the PanTHERIA [27] and EltonTraits 1.0 [28] databases to characterise all of the sampled mammals using a range of traits likely to impact on their suitability as hosts for parasites.

Selected traits were: body mass, which is a key feature of mammals in terms of their metabolism and adaptation to environments; average longevity, litter size and the average number of litters per year as demographic parameters that could be relevant for within-host dynamics of viruses; diet breadth (calculated as a Shannon diversity index based on the proportional use of 10 diet categories as presented in EltonTraits); range area, which we expect to affect the exposure to other mammalian host species; average temperature and average precipitation within a host’s distribution as an indicator of climatic niche; latitudinal centroid of distribution as an indicator of the general habitat and climate within which hosts are occurring across a gradient from tropical to polar biotas; and habitat as multiple binary indicators of whether a species uses 1) forest, 2) open vegetation, and/or 3) artificial/anthropogenic habitats. Information on specific habitat utilisation was compiled from the International Union for the Conservation of Nature (IUCN) database (http://www.iucnredlist.org). We did not include a larger set of ecological traits in our analysis to avoid collinearity issues.

Phylogenetic relationships between sampled mammal species were estimated from a recent mammalian supertree [29]. We used this tree to compute pairwise phylogenetic distances based on a correlation matrix of phylogenetic branch lengths [30] and also a vector of phylogenetic distance to humans for all other mammalian host species. We also quantified pairwise ecological distance between sampled mammal species based on a generalised form of Gower’s distance matrices [31] using weighted variables based on all of the ecological trait variables described above, following methods in [32]. Phylogenetic and ecological distance matrices as well as vectors of trait variables were scaled (dividing by the maximum for each distance matrix), so all distance measures ranged from 0 to one. Data formatting and analyses were conducted in R version 3.4.3 [33] and used the packages *ape* [30] for phylogenetic distance calculations and *ade4* [34] for ecological distance calculations.

### (c) Statistical analysis

The primary focus of this paper was to explore which mammalian host species might be the most important for spreading viruses due to their sharing of viruses with others, and we were interested in the phylogenetic and functional diversity of host species infected by different virus species. We addressed these aims using three different statistical approaches, which we describe in detail in the SI Appendix. In brief, we used the following statistical approaches:

#### Centrality of host species in networks of virus sharing

We calculated eigenvector centrality (a generalization of degree, which is the number of connections a host species has to others in terms of virus sharing; eigenvector centrality accounts both for the degree of a host species and those of connected species, i.e. it considers host species to be highly central if their connected species are connected to many other well-connected species [35]). Eigenvector centrality was strongly correlated with degree measures, betweenness centrality, and closeness centrality (all Spearman r ≥ 0.76). Thus, we present only results from eigenvector centrality and acknowledge that because of collinearity, it is not possible to distinguish further between the different components.

We used the non-parametric Kruskal-Wallis test to assess whether the eigenvector centrality measures differed between wildlife and domestic species and among host orders. We applied Dunn’s test for multiple comparisons [36]. To account for sampling bias that could bias centrality measures [37], we randomly removed subsets of interaction records from the adjacency matrix used for calculating centrality measures. For this, we varied the proportion of removed interactions between 5 – 30% in each of 200 iterations following a uniform distribution. We used the relative proportion of publication and sequence numbers for each mammal-virus combination as two independent sets of probabilities of which interactions to remove. We then calculated centrality measured for each iteration and tested for consistency of results from subsets and the full dataset.

#### Hierarchical model of virus sharing among host species

We generated a binary *N*×*N* adjacency matrix with *z(i,j)* = 1 if the pair of host species *i* and *j* were recorded to share any virus and *z(i,j)* = 0 otherwise (with *i* and *j* ϵ 1,…,*N* and *j* ≠ *i*). The probability *φ(i,j)* that two host species share any virus can be linked to *z(i,j)* with a Bernoulli distribution given as
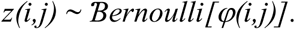
We used the logit-link function to model variation in *φ(i,j)* as

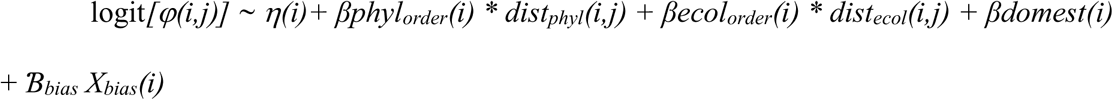

Here, *η(i)* is the species-specific intercept, which is further modelled with a hierarchical hyperprior *η(i)* as ∼ N[H_*η*_*(order)*, σ_*η*_*(order)]*; the hyperprior H_*η*_ accounts for the ‘average’ virus sharing probability of species from different orders, while the variance *σ*_*η*_ accounts for the deviation of species-level virus sharing-probabilities from the respective order-level hyperprior. The coefficients *β*_*phyl*_ and *β*_*ecol*_ account for variation in virus sharing with increasing phylogenetic and ecological distance from *i*. The coefficient *β*_*domest*_ accounts for variation in virus sharing among all possible combinations between species classified as wildlife, domestic, or human compared to pairs of wildlife-wildlife species (a five-level categorical variable), while *μ*_*bias*_ accounts for variation in relation to the four different proxies of sampling efforts described above. We fitted the model in a Bayesian framework with Markov Chain Monte Carlo (MCMC) sampling in the software JAGS version 4.3.0, operated via the R package *rjags* [38].

#### Hierarchical model of the proportion of zoonotic viruses carried by different host species

We modelled the probability *ψ(i)* that a virus recorded for a host species *i* is zoonotic (corresponding to the likely proportion of zoonotic viruses carried by a host species) using a binomial distribution based on the number of zoonotic viruses *y(i)* out of the total number of viruses *w(i)* as
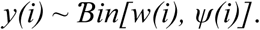
We then used the logit-link function to model variation in *ψ(i)* among different host species as
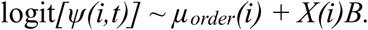
Here, *µ*_*order*_ denotes the order-specific average according to the taxonomic order of species *i*, which were modelled with a Gaussian error structure and a common ‘average’ hyperprior mean, i.e. *µ*_*order*_ ∼ Ɲ(*H*, σ^2^). *X* is a matrix of the 17 species-level covariates (including phylogenetic distance to humans and the four proxies of sampling bias) described above and *B* is a vector of corresponding coefficient estimates. We fitted the model in a Bayesian framework in JAGS [38].

## 3. Results

Of 2,213 virus species associated with 816 different mammalian host species (including humans) in our dataset, 446 species (39%) have been recorded to infect humans. Out of these, 174 species (39% virus species infecting humans) are recorded as zoonotic. Of these zoonotic species, 70 species (40 %) were recorded in wildlife species but not in any domestic species, while 20 species (11 %) were recorded in humans and domestic animals but not in any wildlife species. In turn, 104 (5%) of all recorded virus species were shared by at least one domestic and one wildlife species without being associated with humans.

The virus species included 954 DNA virus species and 1,169 RNA virus species (90 classified as others), of which 42 (4% of DNA virus species) and 119 (10% of RNA virus species) were recorded as zoonotic. The overall network topography for DNA versus RNA viruses reveal distinct spread of these viruses among host species, mostly depicted by considerably lower virus sharing across orders of host species for DNA viruses (**figure 1**).

**Figure 1.**
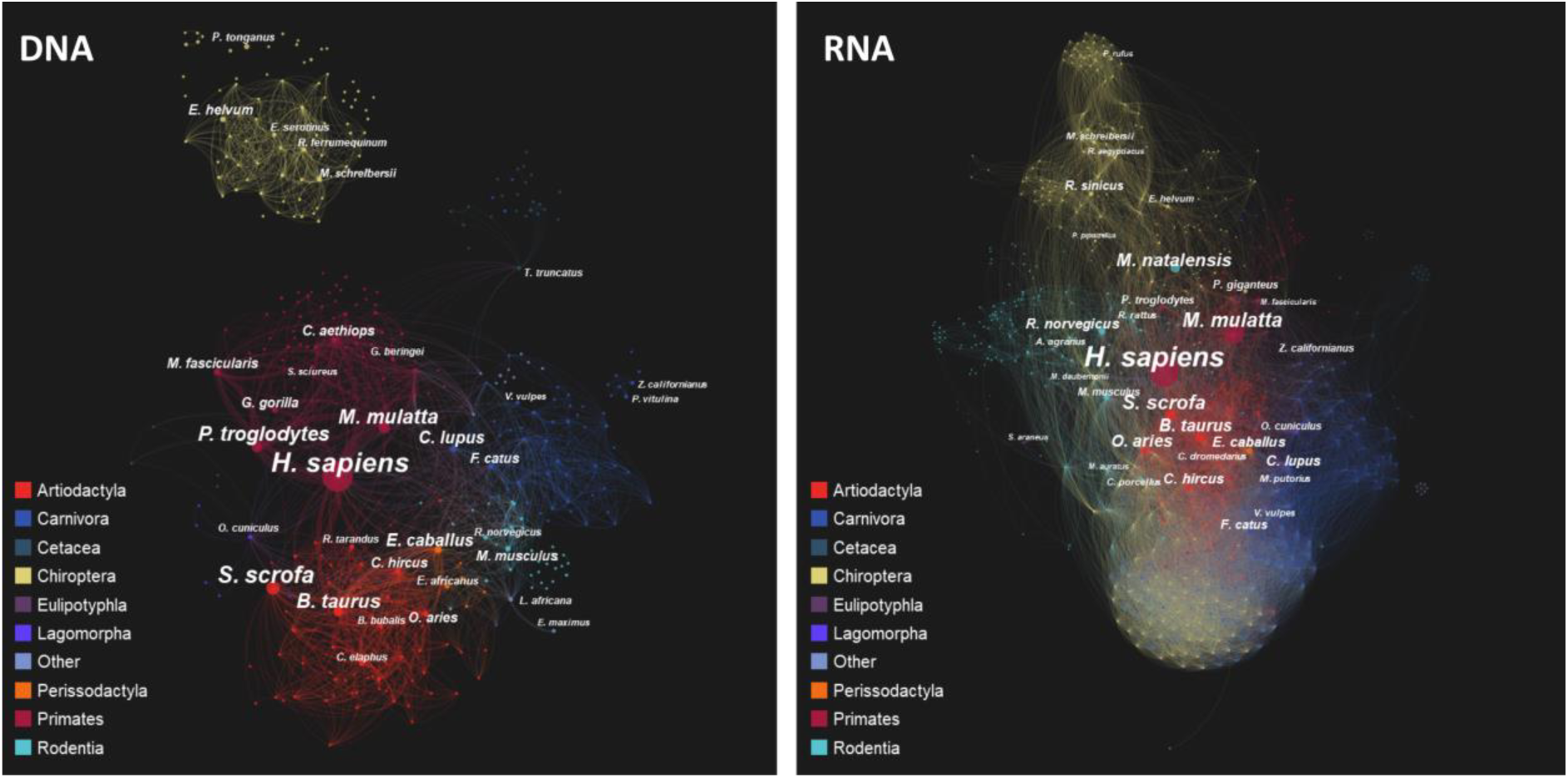
Network plots of the sharing of DNA (left panel) and RNA viruses (right panel) among mammalian host species. Each node represents a mammal species (total of n=816 species) The size of the node depicts the number of virus species shared with other mammalian host species, the width of edges is plotted proportional to the number of virus species shared between pairs of hosts. Color depict the different mammalian orders.

### (a) Centrality of host species in networks of virus sharing and spread

Eigenvector centrality measures were higher for domestic than wildlife host species (Kruskal-Wallis χ^2^ ≥ 36.2, df= 1, p < 0.01), indicating that domestic species were the most central species (after humans and in addition to some primate species) in the entire mammal-virus association network. The ten most central position in the network of all virus species were occupied by *Homo sapiens, Bos taurus, Sus scrofa, Ovis aries, Capra hircus, Macaca mulatta, Canis lupus, Mus musculus, Equus caballus*, and *Pan troglodytes* (following order of descending centrality).

Centrality measures also varied among the different taxonomic orders of host species (all Kruskal-Wallis χ^2^ ≥ 224.7, df = 9, p < 0.01) (**figure 2**). Specifically, eigenvector centrality measures for all virus species were largest for wildlife species of the taxa Carnivora, Chiroptera, Artiodactyla and Primates compared to other taxa (Cetacea, Rodentia, Perissodactyla, others) according to post-hoc multiple comparisons (electronic supplementary material, **table S1**). RNA viruses but *not* DNA viruses accounted for relatively larger centrality scores for Carnivora and Chiroptera (both Mann-Witney U test of group-level comparisons p < 0.01), whereas centrality scores calculated for DNA viruses only were relatively larger for Primates (M.-W. U test p < 0.01) and of equal ranks for Artiodactyla (M.-W. U test p = 0.81) (electronic supplementary material, **figure S1**).

**Figure 2.**
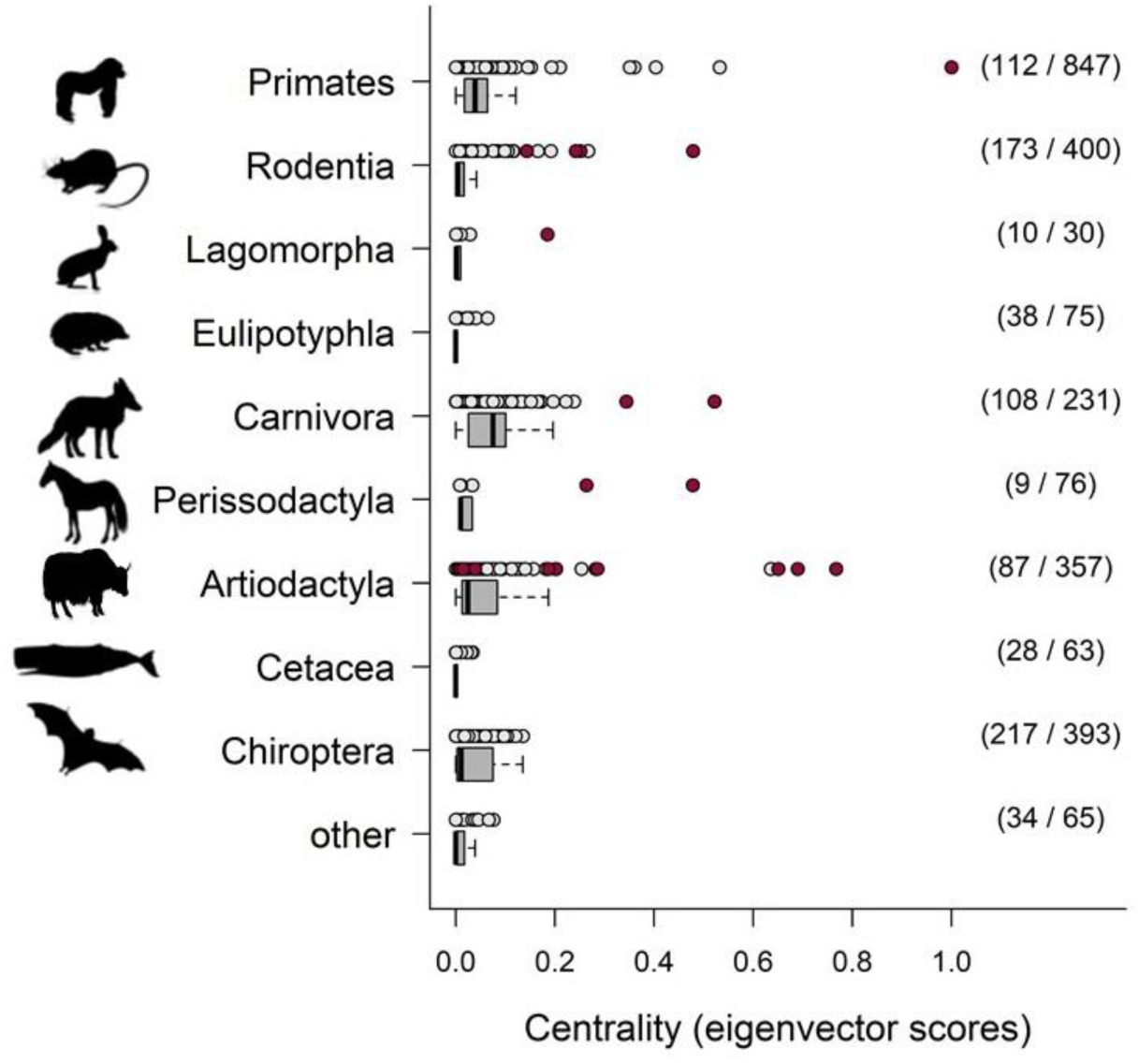
Eigenvector centrality scores (box plots and species data points) of host species from different mammalian orders, depicting their relative importance in virus sharing and spread across the entire network of mammal-virus associations. Larger values refer to host species sharing more viruses with others, especially with host species that are also well connected. Artiodactyla and Cetacea are presented as separate groups because of their distinct ecological niche, mammalian orders with few species are merged into the group ‘other’. Grey points represent measures for wild and red points measures for domestic mammalian host species and humans. The respective number of mammals and associated virus species in each group are given in parenthesis.

Centrality measures calculated from subsets of underpinning adjacency matrices for all viruses with 5 – 30% of interactions removed according to number of publications and published sequences revealed a 4-fold stronger decline in correlations for the number of publications than published sequences (electronic supplementary material, **figure S2**). For these data subsets, there were a total of 33 host species that emerged as the top ten host species according to centrality measures calculated from data subsets (electronic supplementary material, **figure S3**). However, despite this uncertainty in which host species occupied the most central positions, the findings of significant larger centrality measures for domestic than wildlife species hold true for all subsets (all Kruskal-Wallis tests with χ^2^ ≥ 16.5, df= 1, p < 0.01) (electronic supplementary material, **figure S1**).

### (b) Virus sharing among host species

Analysing virus sharing patterns in a probabilistic hierarchical modelling framework confirmed the prominent role of domestic animals in virus sharing across the entire network. Wild mammalian host species were ca. 10 times (95% credible intervals [CIs] of odds ratio 8.7 – 15.5) more likely to share virus species with humans and ca. 6 times (odds ratio 6 – 6.8) more likely to share virus species with domestic animals than with any other wild species. Any pair of domestic species was ca. 84 times (odds ratio: 55 – 141) more likely to share viruses than any pair of two wildlife species. We found the highest frequencies of virus sharing for RNA viruses shared among species of the orders Carnivora and Chiroptera (averaging frequencies of 1.2 – 3.4% according to CIs of sharing RNA viruses with other species), whereas DNA virus sharing frequencies were below 0.7% (according to all upper bounds of CIs) for all host orders (**figure 3**). For all host orders and both virus genome types, we found virus sharing to be more likely with closely related species (negative values for coefficients *β*_*phyl*_ that depict increasing virus sharing for smaller phylogenetic distances among pairs of host species). Phylogenetic clustering of host species (which translates into higher host specificity for the viruses) was stronger for DNA viruses shared by Primates, Lagomorpha, Artiodactyla and Chiroptera compared to RNA viruses (**figure 3**), signifying a general tendency of higher host specificity in terms of phylogenetic similarity for DNA viruses compared to RNA viruses. Notably, phylogenetic host specificity for RNA viruses shared by Primates were relatively low, suggesting host sharing with phylogenetically distantly related host species (**figure 3**). Functional distance among host species were generally less meaningful in describing patterns of virus sharing among pairs of host species, but we found virus sharing for Primates, Rodentia, Artiodactyla and Chiroptera with other hosts to increase with increasing functional distance (positive values for coefficients *β*_*ecol*_ that depict increasing virus sharing for larger functional distances) (**figure 3**). Virus sharing among host species increased with all four proxies of sampling bias for both DNA and RNA viruses (all CIs of odds ratios 1.01 – 2.20), indicating that sampling effort impact the topography of currently known mammal-virus networks.

**Figure 3.**
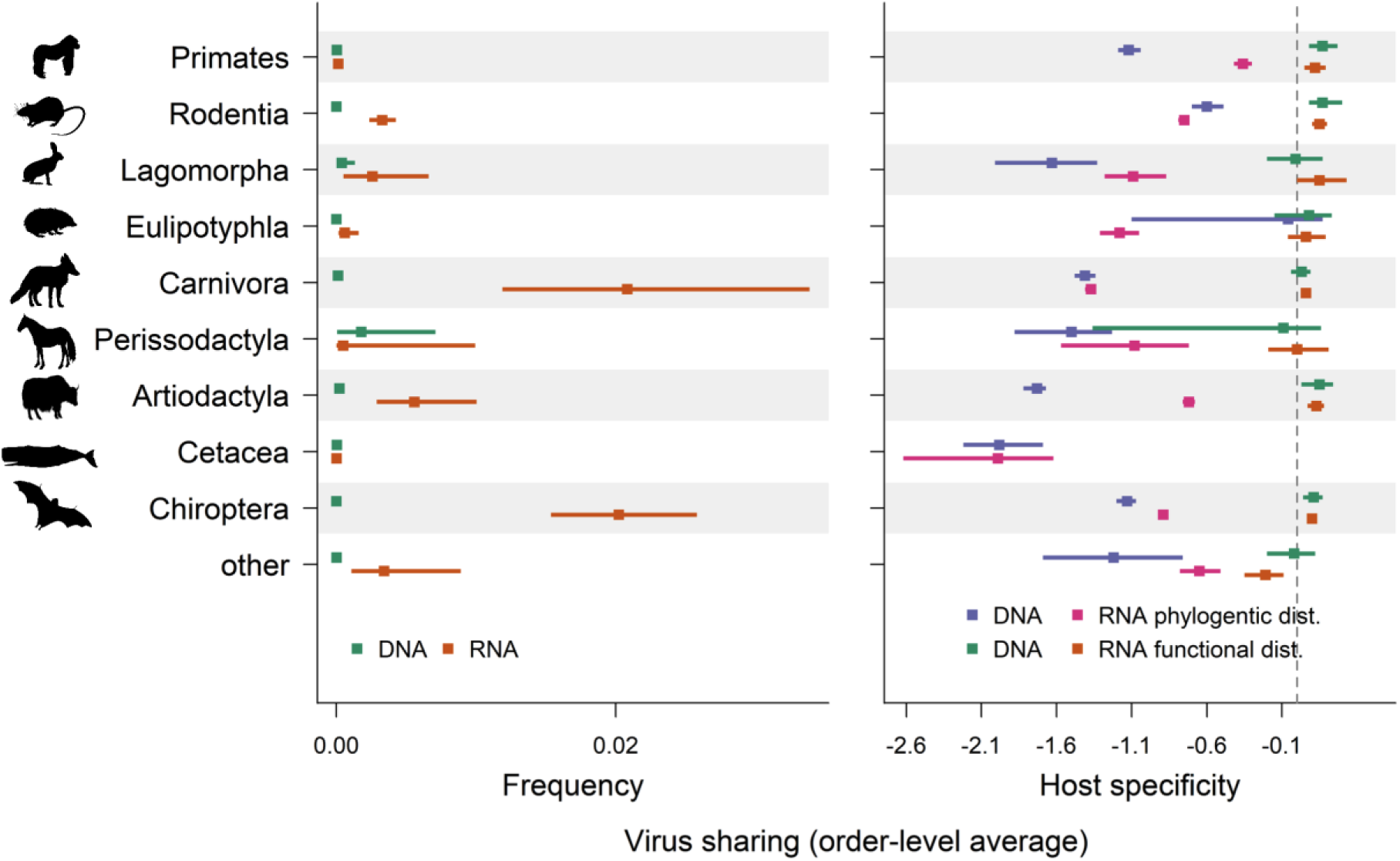
Order-level estimates of the average frequency (left panel; parameter H_*η*_(order) in model description) mammalian species of the respective order share any of its associated viruses with another mammalian host species. The right panel shows the relative extent of host specificity in virus sharing in terms of the relative difference between observed and expected phylogenetic and functional diversity of mammalian host species as estimated from regression coefficients. Values < 0 indicate pairs of infected hosts were more phylogenetically/functionally similar than expected based on random draws from regional mammalian species pools, indicating higher specificity in virus spread among mammalian species (corresponding to parameters *βphyl* and *βecol* in model description). All estimates are presented for the two subsets of DNA and RNA viruses.

### (c) Proportion of zoonotic viruses in different host species

We found the taxa Primates to harbour the overall largest proportions of zoonotic viruses with a group-level average of 54% (CI of 44 – 64% according to *µ*_*order*_) (**figure 4**), followed by slightly lower proportion of zoonotic viruses in Rodentia, Carnivora, Artiodactyla and Chiroptera (all CIs ranging between 14 – 46%) (**figure 4**). The proportion of zoonotic viruses carried by domestic species was 1.2 times higher than in wildlife (odds ratio of 2.2 and CI of 1.5 – 3). RNA virus species accounted for the highest proportions of zoonotic viruses in all mammalian groups, averaging to 46% (CI of 17 – 65% according to hyperprior *H*_*RNA*_) compared to only 7% (CI of 2 – 25% according to hyperprior *H*_*DNA*_) of the DNA viruses in mammalian hosts being zoonotic.

**Figure 4.**
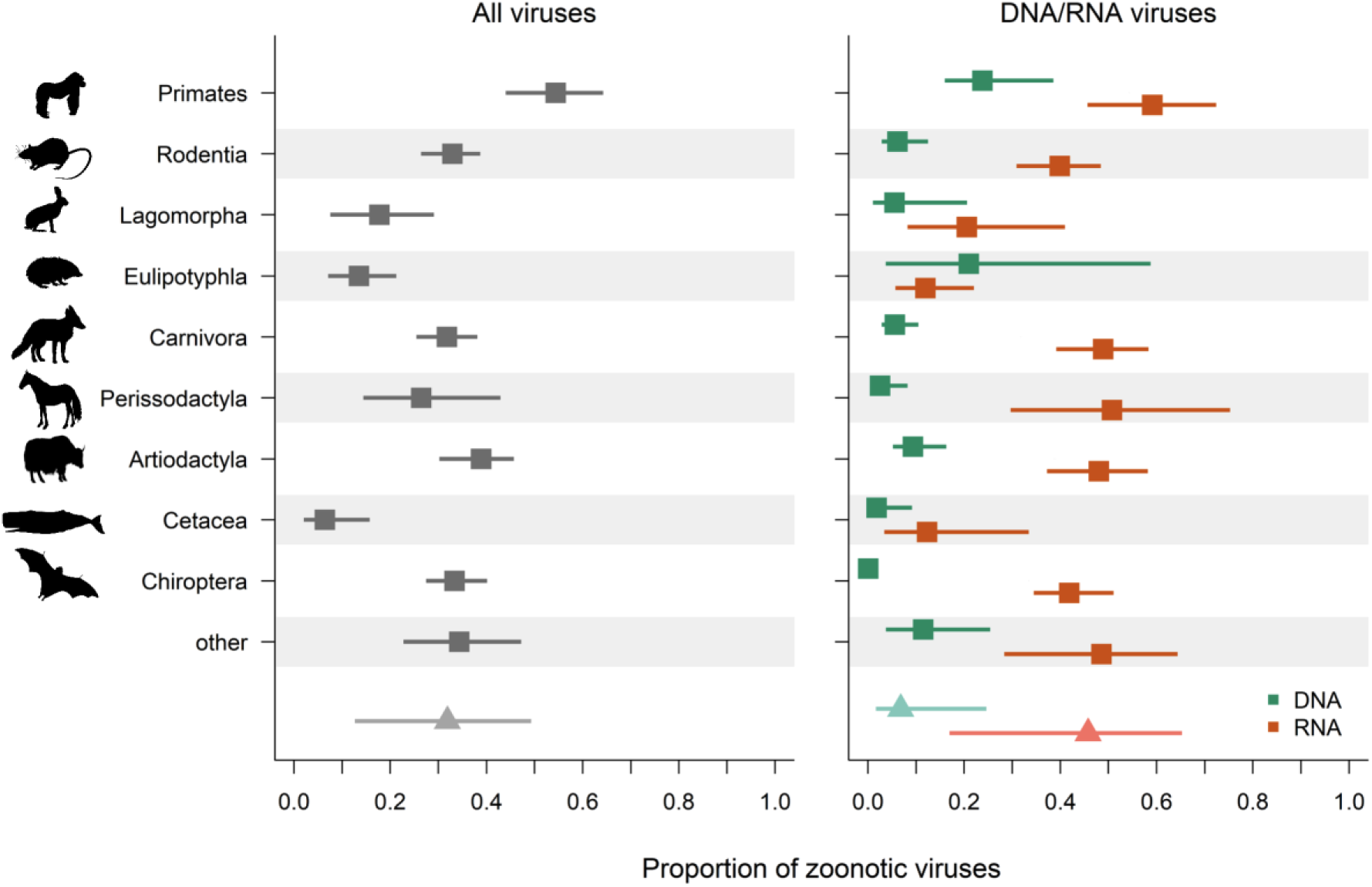
Estimated proportion of zoonotic viruses for mammalian host species from different orders (left panel: all n= 2,213 virus species in the database, right panel: estimates for the two main groups of n=954 DNA virus species and n=1,169 RNA virus species). Estimates represent the group-level averages (‘hyperprior’) from a Bayesian hierarchical model. The group “other” assembles all species from orders with < 9 species in the dataset. Boxes are posterior estimates and bars represent 95% credible intervals. The grey triangle and bar represent the overall average estimate according to a second-level hyperprior in the Bayesian model hierarchy.

We found the proportion of zoonotic RNA viruses in different host species to increase with the numbers of litters per year (odds ratio of 1.1 – 1.6) and larger range area (odds ratio of 1.02 – 1.3), while it decreased with increasing longevity (odds ratio of 0.5 – 0.8) and decreased in open vegetation (odds ratio of 0.5 – 0.99). In contrast, there was no evidence that the proportion of zoonotic DNA viruses in different host species was linked to any species traits (all odds ratio estimates intersecting with 1). The proportion of zoonotic RNA viruses was smaller for host species, with the largest number of sequences recorded and higher Shannon diversity scores of sequence records (both odds ratios between 0.7 – 0.96), suggesting that more intensive sequencing efforts of a large range of these viruses increases the discovery of viruses confined to non-human hosts. The proportion of DNA virus species in different hosts, in turn, increased with higher Shannon diversity scores of publication numbers (odds ratio 1.03 – 1.3).

The associations between host species from different mammalian orders and viruses from different families is illustrated in electronic supplementary material, **figure S4**, data are presented in electronic supplementary material, **table S2**.

## 4. Discussion

Pathogen spillover and the emergence of infectious diseases ultimately depend on how pathogens conquer eco-evolutionary barriers to infect novel hosts [39], but spatiotemporal variation in species interaction and pathogen transmission opportunities are proximately driven by host occurrences and community assembly [40, 41]. It comes therefore as little surprise that globally pervasive mammal groups, such as bats and rodents, are often considered to share as many viruses with humans as do primates, our closest relatives [12, 42, 43]. Our study adds novel insights into viruses spread across mammalian communities. Specifically, we provide the strongest evidence to date that domestic animals are the most central species in mammalian host-virus interaction networks. We also found rather distinctive patterns of how DNA and RNA viruses are shared and spread among different mammalian groups, with bats and carnivores being only of minor role in spreading DNA viruses through the network. We emphasize the dominant role of domestic species in virus sharing, since domestication status strongly increases the chance of virus sharing among multiple mammalian hosts. Likewise, we found domestic species also to carry larger proportions of zoonotic viruses than ‘equivalent’ wildlife species after accounting for phylogeny and other traits.

Our study concerns the contemporary pattern of virus sharing of mammal species rather than any specific co-evolutionary histories of host switching and origin of viruses. In many, perhaps most instances, this sharing indicates the possibility of cross-species transmission, either directly via contact, or indirectly via air, soil, water, fomites or vectors. In some instances, though, the contemporary cross-species transmission of a virus between known host species is no longer possible or is highly unlikely, which is the case if viruses have evolved into distinct lineages that will only spread among closely related host species in response to adaptive evolution such as lineages of rabies viruses confined to bats rather than the entire host spectrum [44]. Some cross-species transmissions may comprise ‘dead-end’ spillovers, if the pathogen is not able to maintain transmission cycles within the novel host [44]. However, the exceptionally high virus sharing of humans and domestic animals with other mammalian species suggest that these species play a crucial role in spreading viruses, as frequent virus acquisition and dissemination is the most plausible explanation for such intensive virus sharing. This may reflect the wide geographic distribution and contact opportunities to wildlife across biogeographic borders, given that domestic species are not particularly distinguished from wildlife in terms of ecological traits. In fact, contact opportunity and community assembly have been shown in a number of studies to impact pathogen sharing and host shifting [8, 45]. Many pathogens, including viruses, can overcome species and environmental barriers to infect distantly related hosts and disperse across large geographic areas [46, 47], although strong constraints in host shifting may also cause biogeographic structure in pathogen diversity and zoonotic disease risk [6, 48]. Beside the large geographic ranges and diverse habitats encroached by domestic species, their large populations sizes and high densities, that often excels those of wildlife populations [18], could further contribute to host shifting and pathogen spread. This could be especially the case if large population sizes facilitate contact opportunity, virus amplification and diversification caused by more intensive within-population transmission or other factors, warranting future research.

Our findings of larger proportions of zoonotic RNA viruses compared to DNA viruses carried in different mammals is consistent with previous research [12, 19, 49] and is in line with our finding that mammal species generally share RNA viruses more frequently with other hosts than DNA viruses. Here, we reveal for the first time that these two major groups of viruses are differently spread across entire networks of mammalian hosts, an important finding that remains largely unnoticed when solely looking at the species richness and propensity of zoonotic viruses carried in different wildlife species. Remarkably, Chiroptera and Carnivora hold central positions in terms of virus sharing with other species only for RNA viruses, while Primates occupy relatively more central positions in terms of DNA virus sharing among hosts. In practice, these findings translate into a minor role of bats and carnivores for the spread of DNA viruses (and relatively low risk that DNA viruses will spillover from these species to humans), whereas at least a few primate species, namely *Macaca mulatta, Pan troglodytes, Chlorocebus aethiops* and *M. fascicularis*, have relatively more central roles in sharing DNA than RNA viruses. We also found that cattle (*Bos taurus*), goat (*Capra hircus*), pig (*Sus scrofa*), horse (*Equus caballus*) and sheep (*Ovis aries*), which are globally the most abundant and economically important livestock species [50], are among those species with the relatively highest centrality measures in terms of DNA virus sharing. Importantly though, it should be noted that for all these species, the frequencies of sharing DNA viruses with other host species was considerably lower than sharing RNA viruses regardless of centrality measures (as is also true for group-level estimates for different mammalian orders as depicted in **figure 3**). We thus emphasize that aforementioned species have a *relative* crucial role in spreading DNA viruses, whereas RNA viruses generally are much more frequently shared among mammalian host species. In this context, our model framework for analysing patterns in host sharing provides probabilistic estimates of the variation in the pairwise phylogenetic and functional similarities of infected versus potential host species as a signal of host specificity. This tool enables us to quantify host specificity of DNA versus RNA viruses in different groups of hosts, resulting in refined and community-wide measures of previously notified higher host specificity in DNA viruses compared to RNA viruses [19-21].

The understanding of virological factors that ensure efficient virus replication and transmission within and among host species is in its infancy [51]. Consequently, disentangling host or virus traits as drivers of the differential spread of DNA and RNA viruses among different mammalian orders is currently not possible and requires fundamental research. Possible working hypotheses as to why primates and ungulates are of relatively high central importance in sharing DNA viruses could be linked to mechanisms that enable efficient within-host virus replication and population-level transmission. At the same time, exploring virus attributes of the major DNA virus families shared among these host species, namely Herpesviridae, Papillomavirridae and Adenovirridae (electronic supplementary material, **figure S4**), may help to explain why these viruses are more likely to be shared by primates and ungulates but are less likely to cross host species barrier with regards to bats and carnivores. Moreover, the strong links of some RNA viruses such as the Bunyavirales to arthropod vectors [52], requires further research into the role of host-vector associations and other transmission modes for the spread of viruses.

We recognize several shortfalls in analysing database records of host-pathogen associations. First, any record of a virus species in a host entirely relies on targeted molecular screening. Certain research foci such as the boost in coronavirus research linked to bats after the SARS pandemics [14] may include a sampling bias difficult to capture when only accounting for publication or sequencing numbers as proxies for sampling bias, since the true presence/absence of viruses in non-target host species remains unknown. Undoubtedly, major research efforts are linked to viruses of public health relevance, while there is a dearth of systematic pathogen surveillances in wildlife [53]. Second, detecting a pathogen in any targeted host species depends on its prevalence in its host population and the number of sampled host individuals but such information cannot be retrieved from database records. With sparse data, any direct interpretation of absolute numbers of species richness and interactions could rather reflect the observation process than true biological patterns and processes [26], and we are therefore currently not able to explore such important properties in our study. Network topologies can be also biased by sampling and data aggregation [54]. We control for research effort in our analysis by accounting for variation in relation to publications and sequencing numbers, as has been done previously [10, 12]. However, as more complete data from systematic disease surveillance efforts becomes available, it will be desirable to improve such analysis to better distinguish true but undiscovered interactions from ‘false zeros’ among other sources of bias. Finally, we are aware that amalgamating species-specific host-pathogen interactions into *N*×*N* adjacency matrix as used for some network statistics comes at the cost of losing information about pathogen species identity and thus overall connectivity of host species can no longer be traced back to particular pathogen species. Overall, network connectivity and modularity are therefore community-level entities, while a focus on particular virus species would require more detailed analysis of underlying species-level interaction matrices.

Our work reveals the importance of domestication status and phylogenetic clustering on the importance of virus sharing among mammals, showcasing also the limited sharing of DNA viruses by bats and carnivores in contrast to primates and ungulates species that readily share both RNA and DNA viruses. The emergence of novel infectious diseases through pathogen spillover is a hierarchical process. Ecological factors that determine the contact opportunity between different host species pave the way for cross-species transmission, host adaptation and subsequent within-host reproduction and transmission, which are then largely controlled by ecophysiological and genetic factors. Future work that better accounts for virus factors and host species community assembly may shed further light on why different types of viruses spread differently among phylogenetic and functional groups of mammals and foster better predictions of future disease emergence.

## Data accessibility statement

The data reported in this paper will be deposited at Dryad (*https://datadryad.org/*).

## Authors’ contributions

M.W. and M.B. established the EID database, M.W. compiled data. K.W. analysed data and wrote the first draft. All authors contributed to interpretation of the results and writing.

## Acknowledgments

We are grateful to reviewers for critical feedback that helped improve an earlier version of the paper.

